# Peak shifts and extinction under sex-specific selection

**DOI:** 10.1101/2021.02.18.431780

**Authors:** Stephen P. De Lisle

## Abstract

A well-known property of sexual selection combined with a cross sex genetic correlation (*r*_mf_), is that it can facilitate a peak shift on the adaptive landscape. How do these diversifying effects of sexual selection + *r*_mf_ balance with the constraints imposed by such sexual antagonism, to affect macroevolution of sexual dimorphism? Here, I extend existing quantitative genetic models of evolution on complex adaptive landscapes. Beyond recovering classical predictions for the conditions promoting a peak shift, I show that when *r*_mf_ is moderate to strong, relatively weak sexual selection is required to induce a peak shift in males only. Increasing the strength of sexual selection leads to a sexually-concordant peak shift, suggesting that macroevolutionary rates of sexual dimorphism may be largely decoupled from the strength of within-population sexual selection. Accounting explicitly for demography further reveals that sex-specific peak shifts may be more likely to be successful than concordant shifts in the face of extinction, especially when natural selection is strong. An overarching conclusion is that macroevolutionary patterns of sexual dimorphism are unlikely to be readily explained by within-population estimates of selection or constraint alone.

## Introduction

A longstanding dilemma in evolutionary biology lies in understanding how populations can evolve from one phenotypic optimum to another. When a population is under net-stabilizing selection and in the vicinity of the optimum trait value (a ‘peak’), selection will pull the population towards the nearby optimum (1), leaving alternative optima seemingly inaccessible (2). For a peak shift to occur, some force must allow a population mean phenotype to transcend the pull of the nearby optimum, and cross a fitness valley to climb a peak beyond (3-6).

Candidate phenomena that may facilitate crossing a natural-selection valley include genetic drift, a change in the environment, or sexual selection. Wright famously proposed a key role for drift in valley crossing (3, 5, 6), although drift alone will only facilitate such a crossing with exceptional rarity, requiring very weak selection (a shallow valley) and small population size (7-9). A change in the environment seems a likely explanation, although also an incomplete one as rates of phenotypic macroevolution do not seem to be coupled to environmental upheaval in any obvious way (10, 11). Finally, sexual selection can readily pull a population off a natural-selection fitness optimum (12), resulting in a peak shift even across quite deep valleys (13). If only one sex is under significant sexual selection, whether or not a peak shift occurs in the other sex depends on the magnitude of the cross sex genetic correlation (*r*_mf_) for the trait (13). If *r*_mf_ is high enough, sexual selection in one sex will pull both sexes off of their optimum, leading to a peak shift in both sexes. More generally, directional selection on any trait can induce a peak shift in other, genetically correlated traits that themselves reside on an optimum (9). In this way, sexual selection coupled with cross-sex genetic correlations has been proposed as a likely mechanism facilitating peak shifts and thus promoting the origin of diversity (14).

Two open questions remain in light of sexual selection’s likely role in driving peak shifts. First, how do sexual selection-induced peak shifts manifest evolution of sexual dimorphism, when adaptive landscapes are complex; That is, when sexually dimorphic and sexually-monomorphic trait optima exist, what are the conditions that promote or constrain likelihood of a peak shift to each optimum? Second, how do we reconcile the diversifying effects of sexual selection and *r*_mf_ with the constraining effects that *r*_mf_ is expected to have on male and female adaptation? Put another way, although we know sexual selection + *r*_mf_ can lead to a peak shift, we also know that this condition of sexual conflict constrains adaptation.

In this note I extend Lande’s (12) model of sexual and natural selection towards a single optimum to the case of multiple optima. This model is similar to one analyzed by Lande and Kirkpatrick (13), but is agnostic to female preference evolution, a major focus of their model. My analysis reveals two underappreciated features of peak shift models. First, large regions of parameter space exist in which relatively weak sexual selection is required to induce a sex-specific peak shift; strong sexual selection is expected to lead to evolution along a line of sexual monomorphism at the macroevolutionary scale. Second, the latter type of peak shift is severely limited by extinction. Although these effects are consistent with results of previous work (9, 13), to my knowledge they have rarely been appreciated, particularly in the context of macroevolution of sexual dimorphism.

### The Model – Two-sex ‘Twin Peaks’

Lande (12) considered a model of evolution of sexual dimorphism by natural and sexual selection, in which natural selection favors a single optimum value for the male trait *z* and female trait *y*. His model of natural selection corresponds to a single adaptive peak on the two-sex adaptive landscape (where 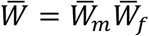, Figure S1A). Directional sexual selection acts independently of natural selection to redistribute fitness across individuals without changing mean absolute fitness. This scenario represents a constant mating bias for one sex, arising from, e.g., female mating preference or male-male competition.

We can expand natural selection fitness function of the Lande model to consider a scenario where more than one optimum for a trait exists, using a mixture of Gaussian functions (15) termed the ‘Twin Peak’ model by Price et al. (9). For males, this leads to the following function relating population mean fitness 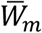 to population mean phenotype 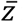:

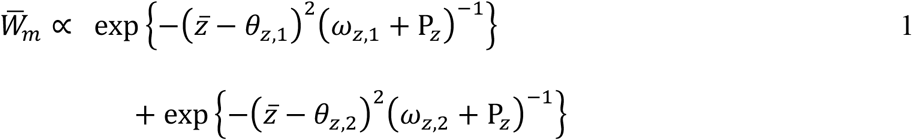

where *θ*_*z*,1_ and *θ*_*z*,2_ are two phenotypic optima, P_*z*_ is the phenotypic variance, and *ω*_*z*,1_*ω*_*z*,2_are the strengths of stabilizing selection (assumed equal throughout). Such an adaptive landscape corresponds to an individual fitness surface (16) with two separate optima. This model is bimodal, with optima in the vicinity of *θ*, for 2*θ*^2^ > *ω* + P. The adaptive landscape for a scenario where only one sex has multiple optima is illustrated in Figure S1B. If instead we assume females also have multiple optima, using an expression analogous to equation 1 but relating 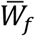 to the female mean trait value 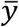, we obtain the ‘Two-sex Twin Peak’ model, illustrated in Figure S1C and Figure 1.

**Figure 1.**
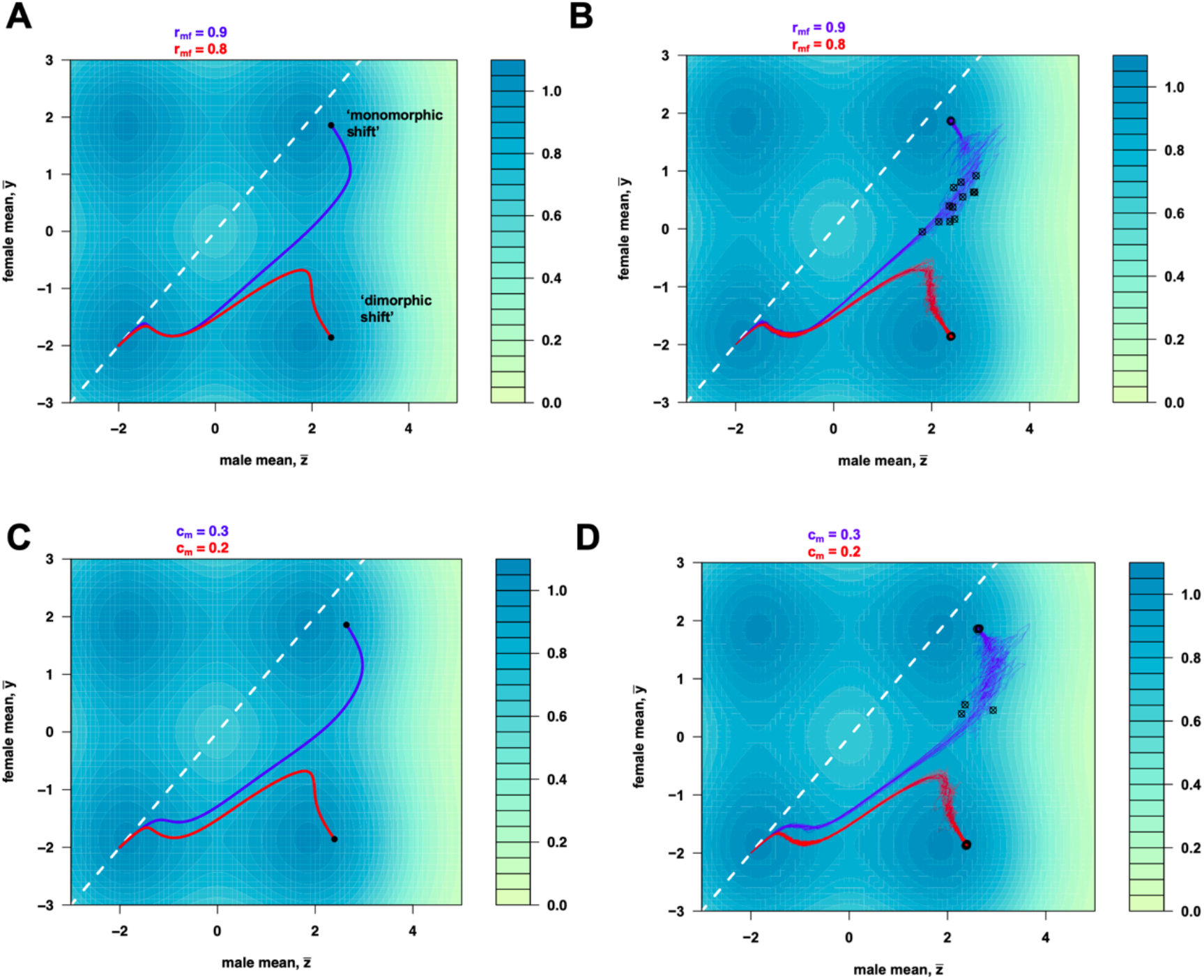
Peak shifts induced by sexual selection in the Two-sex Twin Peaks model. Panel **A** shows two deterministic trajectories, starting from the lower left optimum, corresponding to two different values of the cross-sex genetic correlation *r*_mf_. Although sexual selection is strong enough to induce a peak shift, which new optimum reached depends upon the value of *r*_mf_. Panel **B** illustrates 20 replicates of stochastic evolution (i.e., with drift) under the same parameter values as A and assuming female demographic dominance 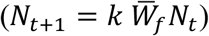. Round circles illustrate the mean values after 10000 generations of evolution; extinction events are denoted with a crossed circle. Panel **C** contrasts trajectories under two strengths of sexual selection c_m_, with otherwise identical parameter values. Panel **D** illustrates stochastic evolution under the same parameter values as C. White dashed line illustrates the line of sexual monomorphism, for reference. In A and B, c_m_ = 0.2; B and C *r*_mf_ = 0.8; other parameter values as described in text.

Given 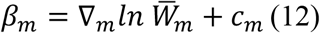, where ∇_*m*_ is the gradient 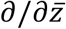 and c_m_ is sexual selection arising from some mating bias independent of the strength of natural selection, we can define *β*_*m*_ under the twin peak model in equation 1 as:

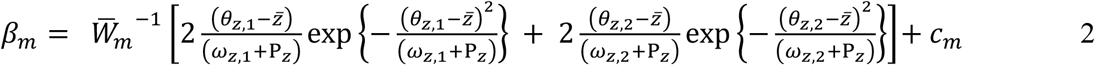

With an analogous expression for *β*_*f*_. For simplicity and consistency with past work, I focus on the scenario of sexual selection in males only (*c*_*f*_ = 0, *c*_*m*_ ≥ 0). Evolution of the male and female mean phenotypes, 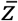 and 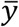, depends not only on selection within each sex but also on correlated response to selection in the other sex mediated by the cross sex genetic correlation *r*_*mf*_:

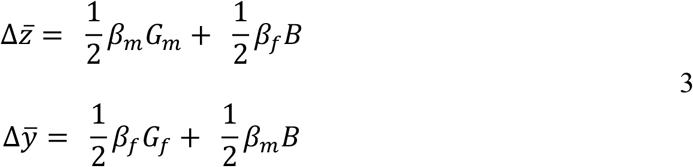

where *G*_*m*_ and *G*_*f*_ are the male and female genetic variances and *r*_*mf*_ = *B*/√(*G*_*m*_*G*_*f*_).

#### Population growth

Population growth can be considered a function of mean fitness, such that maladaptation carries demographic cost. Following Lande (12), we can describe change in census population size N as

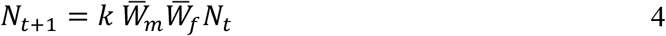

where *k* is constant or function defining per capita birth rates. For simplicity, I assume *k* as a constant corresponding to density-independent population growth. In equation 4, adaptation in both sexes contributes to population growth, corresponding to a biological scenario where, for example, parental care is shared across the sexes. Alternatively, in many species male adaptation may contribute little to population growth rates, and we can instead define change in census size as

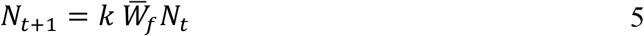

which assumes that there are always at least enough males to fertilize females in the population. Equations 4 and 5 represent two ends of a continuum of in which male adaptation may contribute to population growth rates.

#### Drift

The role of random genetic drift has been considered extensively in peak shift models, and I address it only briefly here for completeness. The rate of evolution by genetic drift will be proportional to **G**/*N*_*e*_ (17), and in a two-sex population,

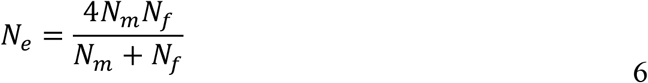

(18). Drift occurs after natural selection such that each sex is represented proportional to their own mean fitness,

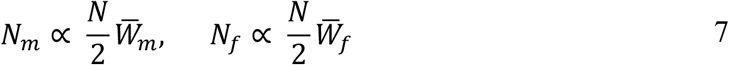

where N represents the number of individual (e.g., zygotes) before selection, with N/2 of each sex under a Fisherian primary sex ratio (1).

I used numerical simulations to explore how sexual selection and *r*_mf_ influence peak shifts and extinction. In order to understand how sexual dimorphism evolves from an ancestral condition of sexual monomorphism, each simulation was started with the population mean at a sexually-monomorphic optimum of 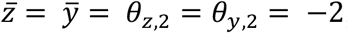, with *θ*_*z*,1_= *θ*_*y*,1_= 2. In this scenario there are three unoccupied optima, two of which are accessible deterministically under positive sexual selection and *r*_*mf*_; one new optimum is along a line of male-female isometry, and represents a shift from one sexually monomorphic peak to another (henceforth, ‘sexually-monomorphic’, or concordant, peak shift). A second accessible peak involves a male-only shift, and so represents a ‘sexually dimorphic’ peak shift. All *ω* were assumed equal, and phenotypic variance was set at unity and G = P/2, assumed constant. Starting N was set to 10,000; population growth was assumed as in equation 3 or 4, although population size was capped at 10,000 with an arbitrary extinction threshold of N = 20. The growth constant *k* was assumed 1.05. Thus growth was exponential and density independent up to the upper bound. Assuming unbounded growth, changing population size, *k*, or genetic parameters did not change qualitative conclusions. Complete R script is provided as supplemental material.

## Results

When there is only a single optimum, sexual and natural selection jointly determine the equilibrium trait value for males, which are displaced from their peak proportional to sexual selection c_m_ and the strength of stabilizing selection *ω* (Figure S1D; 12). Female equilibrium trait values are unaffected by the strength of sexual selection in males, but their path towards their optimum is affected by *r*_*mf*_ (Figure S1D). When multiple optima exist, the combination of sexual selection and *r*_*mf*_ determines if a peak shift occurs and which alternative optima is reached. Two scenarios are shown in Figure 1, illustrating that increasing the strength of sexual selection while holding *r*_*mf*_ constant, and vice versa, have similar effects. Unlike the single optimum case, when multiple optima exist, female equilibrium trait values depend on sexual selection in males and *r*_*mf*_; for a peak shift to occur in females, their product must be high enough for female mean phenotype to be displaced beyond the critical value required for a peak shift. Note that a ‘monomorphic peak’ shift (Figure 1A) still entails some sexual dimorphism in the traits will be observed at equilibrium (because males will be displaced optimum), although the equilibrium magnitude of sexual dimorphism will be far higher when a peak shift is restricted to males only (a ‘dimorphic shift’; Figure 1). Valley crossing carries substantial demographic costs, illustrated in Figure S2.

Whether a peak shift occurs for males depends only on the strength of sexual selection c_m_. However, whether the shift is dimorphic (males only) or monomorphic depends also on *r*_*mf*_, such that for wide ranges of moderate to strong genetic correlations, increasing the strength of sexual selection while holding *r*_*mf*_ constant yields first a transition from no peak shift to a dimorphic shift, and then to a monomorphic shift, and then in some cases extinction (Figure 2, left panels). Increasing the strength of stabilizing selection (smaller *ω*) exacerbates these effects. These effects hold regardless of the demographic model assumed, although extinction is far more likely to be observed when population growth rates are a function of both male and female adaptation (Figure 2, right panels). These conclusions remain qualitatively unchanged when accounting for genetic drift, although the outcomes become probabilistic due to the stochastic nature of drift in the fitness valley (Figure S3).

**Figure 2.**
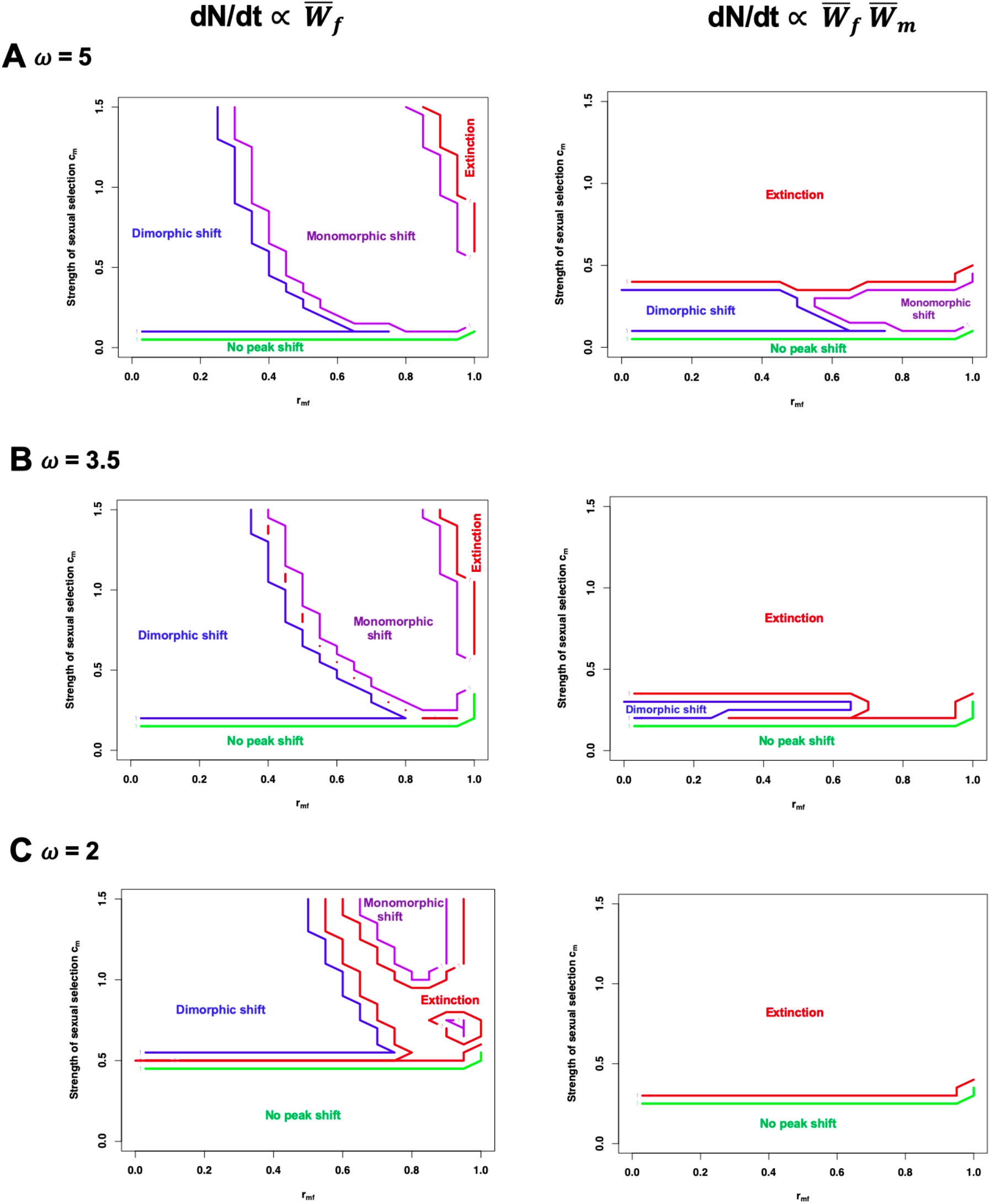
Deterministic outcomes as a function of sexual selection and *r*_mf_. Panels show deterministic (i.e. no drift) outcomes of 1000 generations of evolution, calculated numerically, corresponding to three different strengths of stabilizing selection. Left panels assume female demographic dominance, right panels assume male and female fitness both determine population growth rate. ‘Dimorphic shift’ refers to a male-only peak shift, ‘Monomorphic shift’ refers to a peak shift in both sexes (illustrated in Fig 2A). No peak shift means the lineage remains in the vicinity of the ancestral optimum. Small gaps between regions represent plotting limitations of overlaid contour plots.

## Discussion

Although the broad factors that may possibly affect peak shifts are well understood, how these factors jointly determine the probability of actual peak shifts remains a puzzle (6). Sex-specific selection, in particular sexual selection, provides one example of the uncertainties around peak shifts. Sexual selection and genetic correlations may be key in facilitating peak shifts (12, 13), yet this diversifying effect (14) is seemingly at odds with the constraining role that such a condition must also manifest (12, 19). The point of this paper is to explore how sexual selection and *r*_mf_ interact to influence the likelihood of a peak shift occurring, as well as which alternative optima are most likely to be reached. This modest revisit to classical peak shift models yields two underappreciated phenomenon that have important implications for the evolution of sexual dimorphism.

First, although increasing the strength of sexual selection increases the likelihood of a peak shift, whether that peak shift results in substantial sexual dimorphism depends somewhat counter intuitively on the magnitude of sexual selection. For wide ranges of *r*_mf_, weaker sexual selection produces a peak shift in males only; increasing sexual selection further increases the correlated response in females to the point where females also shift peaks (see also figure 4A in Price et al. 1993). The implication is that when sexually monomorphic optima exist, strong sexual selection is expected to lead to peak shifts along a line of male-female isometry when *r*_mf_ is non-zero. Only when sexual selection is weak enough in magnitude that its interaction with *r*_mf_ fails to push females beyond the critical displacement required for a peak shift, yet strong enough to directly push males beyond their own critical displacement, will dimorphic optima be reached. This suggests that macroevolutionary patterns of sexual dimorphism may depend far more on nuances of the two-sex adaptive landscape than on the magnitude of within-population sexual selection.

Second, the demographic cost of valley crossing affects which peaks can be reached. When population growth rates depend on adaptation in both sexes, or when natural selection is strong, sex-specific peak shifts are more likely to occur successfully in the face of extinction than are concordant (‘monomorphic’) shifts. This demographic effect is the result of higher demographic costs for crossing a valley in both sexes. When population growth rates depend only on female adaptation, valley crossing in males-only carries no demographic cost, and so may be more likely to be observed. These effects are amplified under strong selection (deep valleys), and indicate that extinction-generated survivorship bias (20) may make sex-specific (dimorphic) peak shifts more likely to be observed than monomorphic shifts.

These two features of sex-specific peak shifts may explain several puzzling phenomena observed in macroevolutionary studies of sexual dimorphism. First, a large number of studies have investigated the link between proxies for the strength of within-population sexual selection and among-lineage patterns of sexual dimorphism. Often, these studies find only weak relationships between these sexual-selection proxies and the magnitude of sexual dimorphism (21-24). This finding of little correlation between the magnitude (yet some correlation between the sign; 25) of sexual dimorphism and sexual selection proxies is consistent with the results presented here. Concomitantly, any association between *r*_mf_ and the strength of sexual selection within populations (as some data implies could be the case; 26, 27) could have important macroevolutionary consequences.

Further, when male adaptation contributes little to population growth rate, this may lead to a male bias in macroevolutionary rate if male valleys are easier to cross without extinction. In many clades, males are observed to exhibit higher rates of body size evolution than females (28-31). Although the opposite (higher female rates) is observed in some groups (32), it does appear to be rarer.

Evolution on real adaptive landscapes, with mixtures of peaks of varying height, varying distance from each other, will surely be more complex than the model presented here. None the less, whether a peak shift to a dimorphic or monomorphic peak occurs will depend in part on an interaction between sexual selection and genetic correlations that may decouple the strength of sexual selection from the magnitude of resulting sexual dimorphism at equilibrium, and crossing two valleys at once may be demographically costly.

## Acknowledgements

I thank David Punzalan, Locke Rowe, Erik Svensson, Beatriz Willink, Masahito Tsuboi, and Benjamin Jarrett for discussion and/or comments on the manuscript. Funding was provided by grants from the Swedish Research Council (VR registration number 2019-03706) and Kungl. Fysiografiska Sällskapet i Lund to S.P.D.

## Supplemental Figures

**Figure S1.**
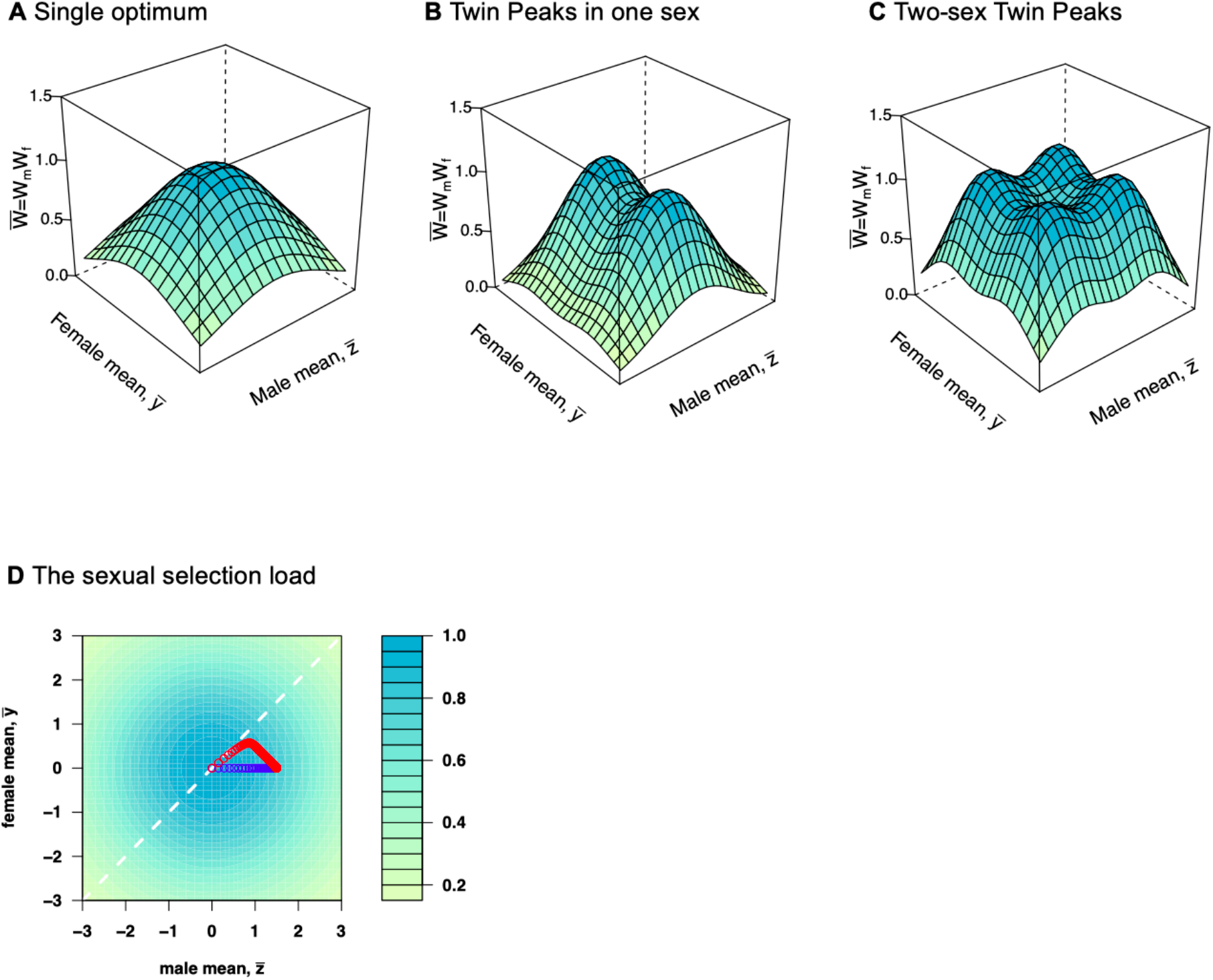
Models of stabilizing selection in two sexes. Panels illustrate the adaptive landscape, the function 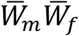. In Lande’s (1980) original model of evolution of sexual dimorphism by natural and sexual selection, a single multivariate optimum, corresponding to one optimum for the male trait and one for the female trait, is assumed (**A**). The ‘Twin Peaks’ model extends this to consider two optima in one trait (sex), with only a single optimum for the other trait (sex); (**B**). Extending the Twin Peak model to both sexes results in two optima for each trait, or four multivariate optima on the two-sex adaptive landscape (**C**). As parameterized in this manuscript, the Two-sex Twin Peak model contains two sexually monomorphic optima, and two sexually dimorphic optima. Of interest is when and how peak shifts from an ancestral monomorphic optimum occur. In this model sexual selection leads to a displacement of males from their optimum (**C**), the magnitude of which is proportional to the strength of stabilizing natural selection. Two trajectories are shown starting from the optimum; one where *r*_mf_ = 0 (blue) and one where *r*_mf_ = 0.9 (blue).

**Figure S2.**
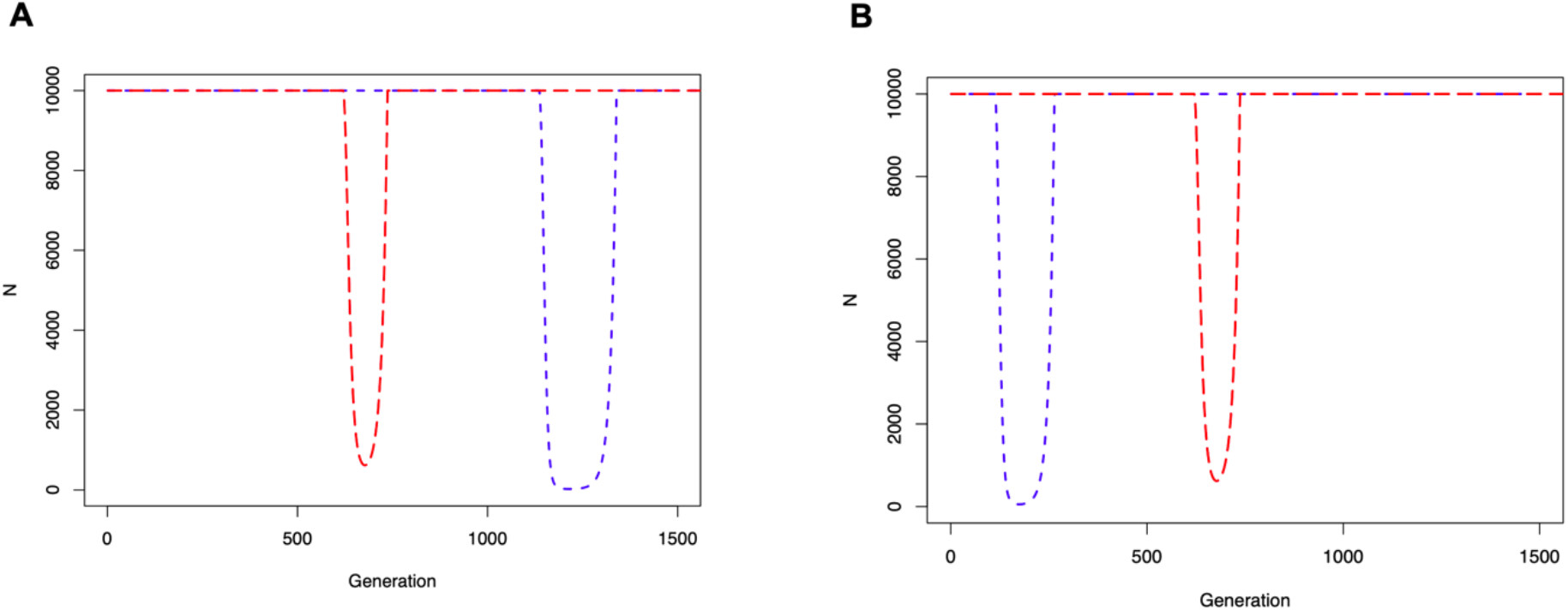
Demographic cost of valley crossing. Figures show population size from trajectories plotted in Figure 1A and 1C. Panel A contrasts two values of the cross-sex genetic correlation, *r*_mf_ = 0.8 (red line) or *r*_mf_ = 0.9 (blue line). Panel B contrasts two values of the strength of sexual selection, cm = 0.2 (red line) or cm = 0.3 (blue line). Demographic costs of a monomorphic peak shift, corresponding to the blue line in both panels, are higher than for a dimorphic (male-only peak shift). Other parameter values are as described in Figure 1.

**Figure S3.**
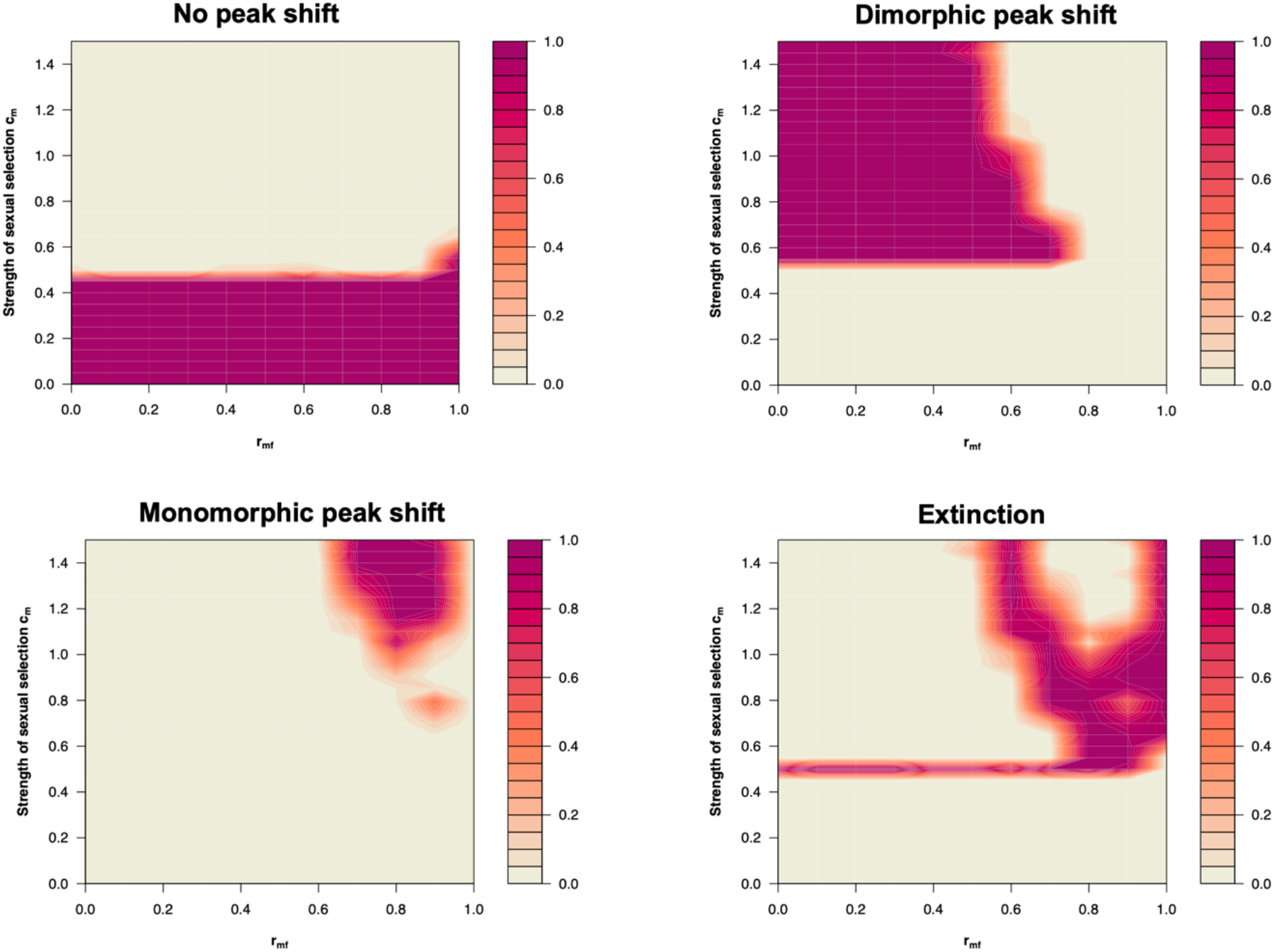
Stochastic outcomes as a function of sexual selection and *r*_mf_. Panels show stochastic (i.e., evolution under both selection and drift) outcomes of 1000 generations of evolution, calculated numerically, corresponding to the scenario of strong natural selection and female demographic dominance illustrated in Figure 2C. Thus, qualitative conclusions on peak shifts remain similar to the deterministic case, and this holds for other parameter values shown in Figure 2.

